# Characterizing Gene Regulatory Network Ensembles in Kidney Injury and Repair

**DOI:** 10.1101/2025.01.12.632586

**Authors:** Eitan Tannenbaum, Dana Markiewitz, Tomer Kalisky, Hillel Kugler

## Abstract

The inference of gene regulatory networks (GRNs) from single-cell RNAseq data allows for mechanistic characterization of the different cell states and their dynamics in complex biological processes. While numerous algorithms have been proposed to infer GRNs from single-cell transcriptomic data, multiple network solutions may explain the same dataset, posing a challenge for biologically meaningful interpretation. Here, we use the Reasoning Engine for Interaction Networks (RE:IN), a computational tool based on formal reasoning, to characterize GRN ensembles in the context of acute kidney injury (AKI). To this end, we applied RE:IN to a single-cell RNAseq dataset from a mouse ischemiareperfusion injury (IRI) model, focusing on distinct proximal tubule cell states related to kidney injury and repair. We first created an Abstract Boolean Network (ABN) model for the kidney using RE:IN and synthesized an ensemble of consistent network solutions. Then, we visualized the ensemble in latent space using Principal Components Analysis (PCA) and discovered four distinct GRN families compatible with the input gene expression and regulatory constraints. Finally, we identified two specific network substructures that discern between the four different network families. This study provides a methodology for characterizing and interpreting GRN heterogeneity in complex processes such as tissue development, disease, and repair.

## 1 Introduction

Gene regulatory networks (GRNs) control the dynamics of gene expression and determine cellular decision-making. These networks can be modeled as graphs consisting of vertices representing transcription factors (TFs) and target genes (TGs), as well as edges describing regulatory interactions [5, 2, 14]. In recent years, algorithms for GRN inference have been developed [1, 9, 17]. However, multiple networks may be consistent with a set of gene expression measurements, and it is a major challenge to characterize and deduce biologically mechanistic meaningful conclusions from such GRN ensembles. In this study we set to characterize GRN ensembles inferred using the Reasoning Engine For Interaction Networks(RE:IN) tool-set [18, 19], a computational modeling tool based on automated formal reasoning. As a biological model system we selected a mouse model for acute kidney injury.

The mammalian kidney is a blood-filtering organ which also maintains blood pressure and acid-base balance. Each kidney contains approximately one million functional units called nephrons. During acute kidney injury (AKI), the kidney experiences structural and functional damage due to ischemic or toxic insults, resulting in loss of epithelial cell integrity, reduced filtration capacity, and disruption of normal cellular processes. Typically, the kidney has the capacity for self-repair. However, in cases of prolonged injury, the epithelial nephron cells fail to repair, driving inflammation and fibrosis, eventually resulting in kidney dysfunction, a situation referred to as chronic kidney disease (CKD). To date, the regulatory mechanisms that determine whether the kidney will heal or develop chronic kidney disease are not well understood. Thus, understanding the GRN governing acute kidney injury is crucial for developing targeted interventions that could prevent the progression from acute kidney injury to chronic kidney disease (CKD).

In recent years, the regulatory interactions governing kidney injury were studied using single-cell RNA sequencing (RNA-seq) and chromatin accessibility (ATAC-seq) [8, 11, 15, 13, 12]. In one study, Kirita et al. [11] used scRNA-seq to examine cell states in a mouse model of bilateral ischemia-reperfusion injury (IRI) over a period from 4 hours to 6 weeks post-injury. They identified several distinct proximal tubule cell states, and labeled them as “severe injured”, “injured”, “repairing”, “healthy”, and “failed repair”. Using SCENIC [1] and GWAS, they identified key transcription factors, target genes, and regulatory interactions. However, it is likely that there are additional GRN topologies consistent with this data. Therefore, we used the RE:IN methodology to investigate the ensemble of GRN topologies.

## 2 Methods

### 2.1 The RE:IN Method

The Reasoning Engine for Interaction Networks (RE:IN) is a computational approach and tool-set enabling automatic construction of GRNs that are consistent with known experimental measurements and known constraints. RE:IN describes GRNs using the Boolean network formalism [10]. In Boolean networks each component can be either On (active) or Off (inactive). The dynamics of the GRN allows the state of the components to progress in discrete time intervals by defining a Boolean variable for each network component and specifying logical update functions according to the interactions and cis-regulatory logic specified by the GRN. According to traditional methods, a Boolean network is manually constructed by the modeler and then compared to known experimental results to ensure consistency with the given data, before being used to make predictions under new conditions. However, traditional methods ignore knowledge gaps and assume a specific structure and cis-regulatory logic to be correct, thus examining only a single network while overlooking alternative solutions. Several approaches and tools, including RE:IN [4, 3, 18], have been developed to mitigate this issue.

RE:IN allows making predictions and analyzing the behavior and structure of biological systems, which are partially known, from a perspective that would have been hard to achieve otherwise. This is achieved by describing systems as Abstract Boolean Networks (ABNs), which are collections of Boolean networks consistent with results, that differ from one another in their update functions for each component. These functions are determined by the interactions that differ between models, which are determined by a set of definite interactions and a set of optional interactions, such that each Boolean network has all definite interactions and a choice of optional interactions. General update functions called regulation conditions (RCs) are assigned to each component and vary across the networks of the ABN [18]. ABNs may then be analyzed for the satisfiability of certain specific behaviors or properties, by performing a process similar to bounded model checking and utilizing an SMT solver [6] to check which models of the ABN satisfy the required behaviors. To be more exact, RE:IN allows to define experimental constraints that will be satisfied by all solutions. These in turn are built of two parts:

1. A Boolean formula called a constraint expression, built of atoms called observations, which specify restrictions on the states of components over time points during multiple executions of our models, corresponding oneto-one to experiment labels specified in the observations.
2. Optional perturbations over certain model components, specifying that these components are inactive or active (knocked out (KO) or forcefully expressed (FE), respectively) indefinitely for certain executions [18].

With that, an ABN, or RE:IN model, satisfies certain constraints if and only if there exists a model and executions corresponding to experiments that make the constraint expression true [18]. The number of models of the ABN that satisfy the constraints is called the number of solutions of the ABN.

### 2.2 Single Nucleus RNA-Seq Data and Preprocessing

A total of six gene expression matrices and their metadata were downloaded from the Gene Expression Omnibus (accession number GSE139107). Only cell identities related to the proximal tubule were selected: “PTS1”,”PTS2”,”PTS3”, “NewPT1” and “NewPT2”. Downstream analysis such as normalization, scaling, and clustering was performed using Seurat v3 [16]. We identified the cell subpopulations shown in Figure 2A of Kirita et al. [11] by checking the expression levels of the specific marker genes shown therein, and labeled them accordingly as “Healthy S1”, “Healthy S2”, “Healthy S3”, “Repairing PT”, “Injured S1/S2”, “Injured S3”, “Severe injured PT”, and “Failed repair PT”.

### 2.3 Pseudo-bulk Preparation and Binarization

The total counts for each gene in each cell type was averaged to create a pseudobulk expression profile for each cell subpopulation. Since the RE:IN algorithm accepts binary inputs (ON or OFF states of genes), we defined the average expression across all cell states as a threshold. The value of each gene was set to 1 if it was higher than the average, or 0 if lower.

### 2.4 Network Clustering and Visualization

Hierarchical clustering was performed using the “ComplexHeatmap” R package [7]. Principal Component Analysis (PCA) was perfomed using the prcomp function of the “stats” R package with default parameters.

### 2.5 Icons

Some icons in Figure 1 were downloaded from Bioicons (https://bioicons.com/). Specifically: (1) the kidney icon was created by Servier (https://smart.servier.com/, licensed under CC-BY 3.0 Unported https://creativecommons.org/licenses/by/3.0/);(2)the singlecell droplet icon was created by Xi-Chen (https://notarocketscientist.xyz); and (3) the Single Cell Clustering UMAP icon was created by James Lloyd (https://www.badgrammargoodsyntax.com/).

**Figure 1:**
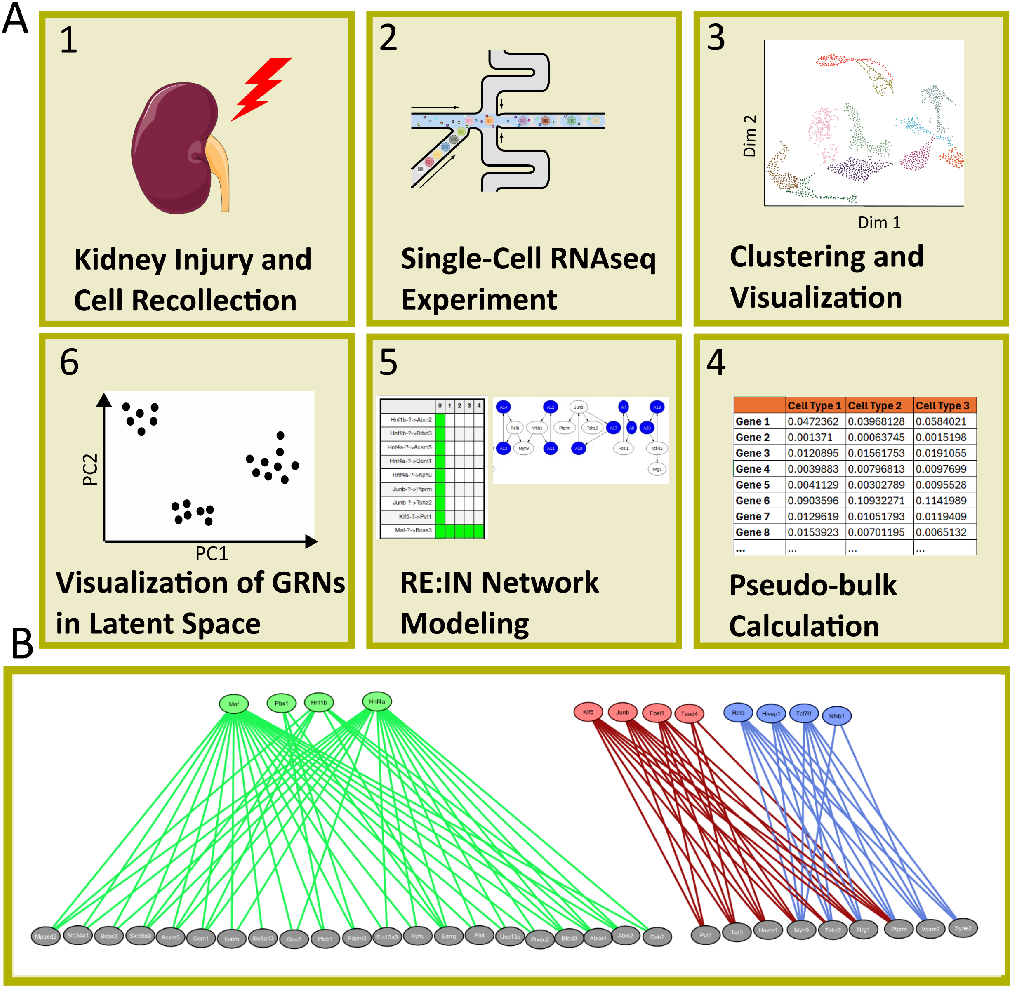
An illustration of the GRN inference and characterization methodology. (A) To infer GRNs controlling kidney injury and repair, cells are collected from a kidney injury experiment and gene expression is measured using scRNA-seq. Then, pseudo-bulk gene expression for each cell state is calculated and used as input to RE:IN for network inference, along with other regulatory constraints. The GRNs are then represented as vectors and visualized in latent space. (B) Constraints imposed on allowed regulatory interactions include those depicted in Figure 3A of Kirita et al. Additionally, the transcription factor Maf was allowed to connect to all target genes regulated by TFs associated with successful repair (green), the transcription factors Klf5 and Junb were allowed to connect to all target genes regulated by TFs associated with injury (red), and the transcription factors Hivep1 and Tcf7l1 were allowed to connect to all target genes regulated by TFs associated with failed repair (blue). Additionally, the target gene Havcr1, associated with injury, was allowed to connect to all related TFs (red), and the target gene Vcam1, associated with failed repair, was allowed to connect to all related TFs (blue).

## 3 Results

### 3.1 Creating the kidney ABN using RE:IN

Our Abstract Boolean Network (ABNs) consists of a set of genes with regulation conditions and interactions between those genes. We focused on 12 central transcription factors previously found to be related to kidney injury and repair [11], and constrained our GRN model to the putative regulatory links showed therein (Figure 1B, see also Kirita et al. Figure 3B). Likewise, we imposed constraints on gene expression dynamics as follows: We downloaded the gene expression matrices of a single-cell RNA-seq dataset collected by Kirita et al. [11] with samples ranging from 4 hours to 6 weeks following IRI, as well as a normal control. We then identified the different cell types (“Healthy S1”, “Healthy S2”, “Healthy S3”, “Repairing PT”, “Injured S1/S2”, “Injured S3”, “Severe injured PT”, and “Failed repair PT”), and created an averaged gene expression profile for each cell type to obtain a pseudo-bulk gene expression matrix. It is thought that the cells in the kidney after injury undergo phenotypic changes resulting in a transition from a severe injured state to a repairing state, after which the cells either repair successfully and transition to their original healthy state, or fail to recover and transition to a “failed repair” state. Therefore, we fitted a RE:IN model for the following trajectory: Severe Injury→Injured S1/2→Repairing (REP)→Healthy S1 or Failed Repair (FR). Since the RE:IN model requires a binary input (ON/OFF) we defined the expression of each gene to be ON or OFF in a specific cell state if its expression was over or under the average expression level. All interactions were assumed to be positive optional. In order to create satisfiable models using RE:IN, a positive time delay was added to each transcription factor by adding two auxillary genes to the network. We also examined other chains of the pseudotime trajectory of proximal tubular cells (Severe Injury (SI) →Injured S1/S2 → Failed Repair (FR); SI→Injured S3 → FR; SI→Injured S1/S2→ Repairing (REP) →Healthy S1; SI→Injured S1/S2→ REP→Healthy S2; SI→Injured S3→ REP →Healthy S3; SI→Injured S1/2→ REP →Healthy S2/FR; SI→Injured S1/2→ REP →Healthy S3/FR) and RE:IN was able to find consistent solutions in these pathways as well.

To summarize, we constructed a network of 41 genes and tested for satisfiability under imposed constraints. The constraints are the values of those genes at designated cell states corresponding to points along pseudotime. We also enabled more flexibility to allow for satisfiability to be achieved. Each transition yielded results. We investigated a solution ensemble of size N=250, 500, 1000, 4000, and 8000. Despite the large state space, RE:IN was able to solve up to 8000 solutions in a matter of several hours, which is a relatively short amount of time for a network of this magnitude.

### 3.2 RE:IN reveals four GRN families compatible with input gene expression and regulatory constraints

The network topology of each RE:IN solution was represented as a vector, where each component corresponds to a specific TF→TG regulatory interaction, and is either one if the interaction exists or zero if not (Figure 2). These vectors were combined into matrices and represented as heatmaps in order to visualize the ensemble of solutions, where each column represents a specific network. Subsequently, we applied Principal Components Analysis (PCA) to project the network solutions onto latent space (Figures 2-3), where each data point corresponds to a network. We observed four discrete well-separated clusters of solutions (Figure 2D,E,F), a division that was robust for N = 500, 1000, 4000, and 8000. This suggests that there are four families of GRNs that are consistent with the observed gene expression and constraints.

**Figure 2:**
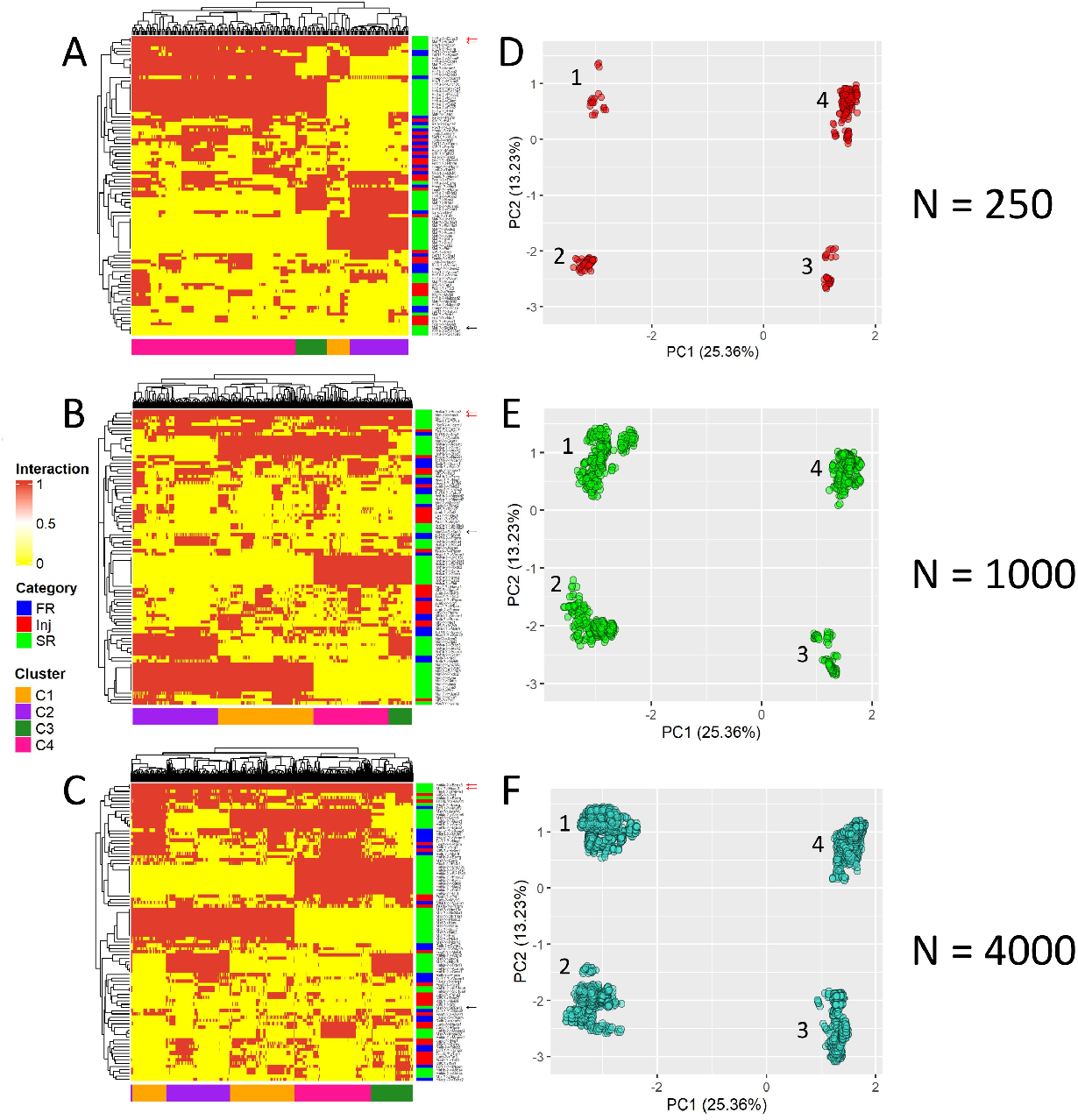
Four GRN families were found to be compatible with input gene expression and regulatory constraints. (A - C) Shown are heatmaps for N=250, 1000, and 4000 solutions. Each column represents a specific GRN solution. Each row represents a TF *→* TG regulatory interaction (Red - exists, yellow - does not exist). Rows and columns were hierarchically clustered with complete linkage and Euclidean distance. GRNs from each of the four families cluster close to each other. Additionally, it can be seen that the regulatory interactions Hnf4a *→* Bcas3 and Maf *→* Bcas3 are required (red arrows), whereas the interaction Maf *→* Slc6a13 is disallowed (black arrow). (D - F) Principal Components Analysis (PCA) shows that the GRNs cluster into four families, and that this division is consistent across N=250, 1000, and 4000 solutions.

### 3.3 Specific network substructures discern between the different four network families

To understand the topological differences between the four network families, we drew representative networks for each of the four clusters (Figure 3), and identified differences in specific network substructures (Figure 4). For example, we found that the transcription factors Maf and Hnf4a switch their target genes (Figure 4A-B), whereby in clusters 1-2 Maf connects to the majority of target genes, whereas in clusters 3-4, the network is rewired such that Hnf4a connects to those targets. The difference between clusters 1 and 4 and clusters 2 and 3, can be explained by alternative linking of specific target genes to the transcription factors Maf, Hnf4a and Hnf1b (Figure 4C-D). In one configuration, Maf and Hnf4a both typically regulate the target genes Acsm5 and Gcnt1, while Hnf1b typically regulates the target genes Btbd3 and Atxn2. In the alternative configuration, Maf and Hnf4a typically regulate Btbd3 and Atxn2, whereas Hnf1b typically regulates the targets Acsm5 and Gcnt1. These switching phenomena can also be observed in the heatmaps (Figure 2).

**Figure 3:**
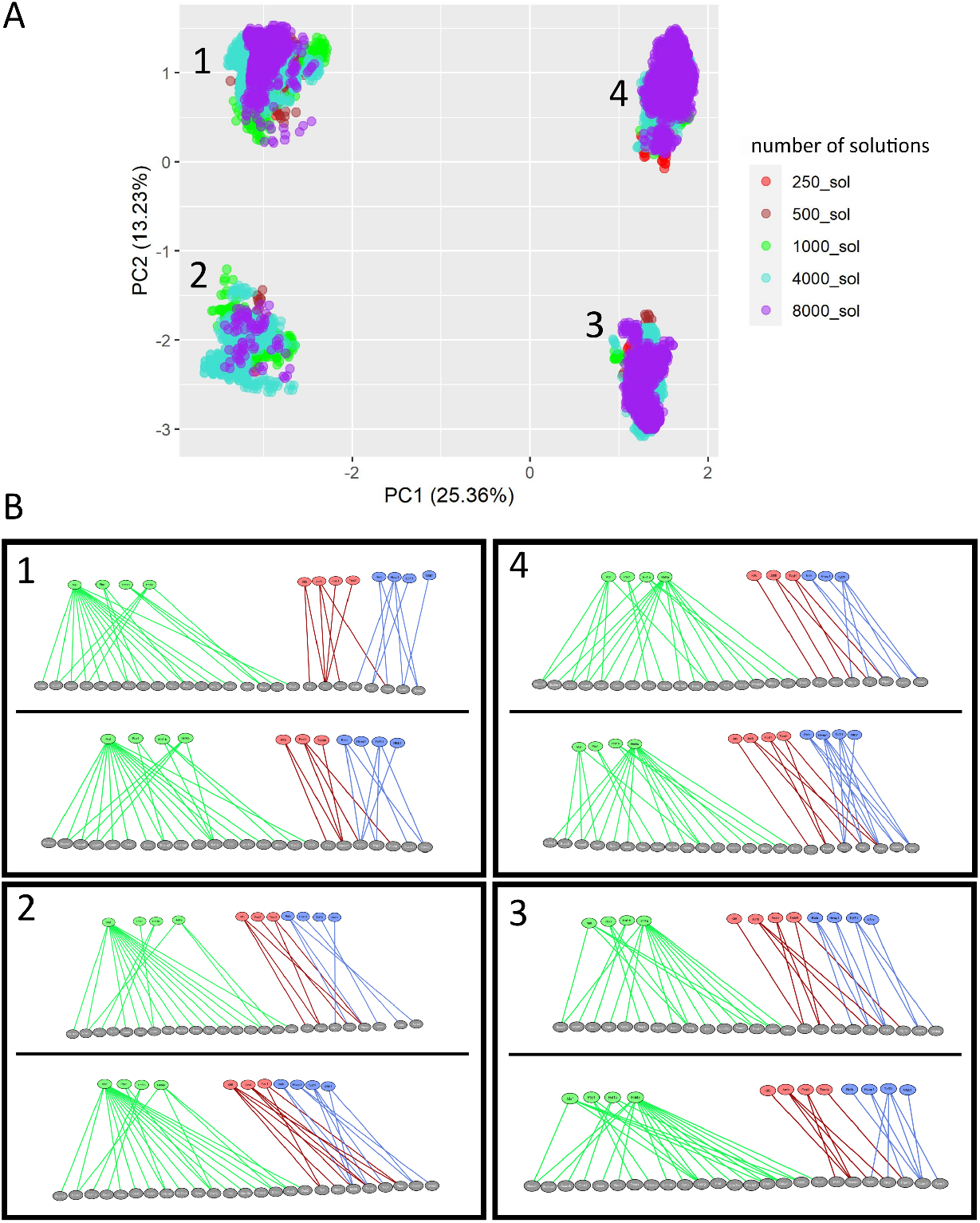
Illustration of specific network representatives from the four network families. (A) PCA shows that the division into four GRN families is consistent across experiments with N = 250, 500, 1000, 4000, and 8000 GRN solutions. (B) Illustrations of representative network configurations are provided for each of the four families, with two GRNs depicted per family.

**Figure 4:**
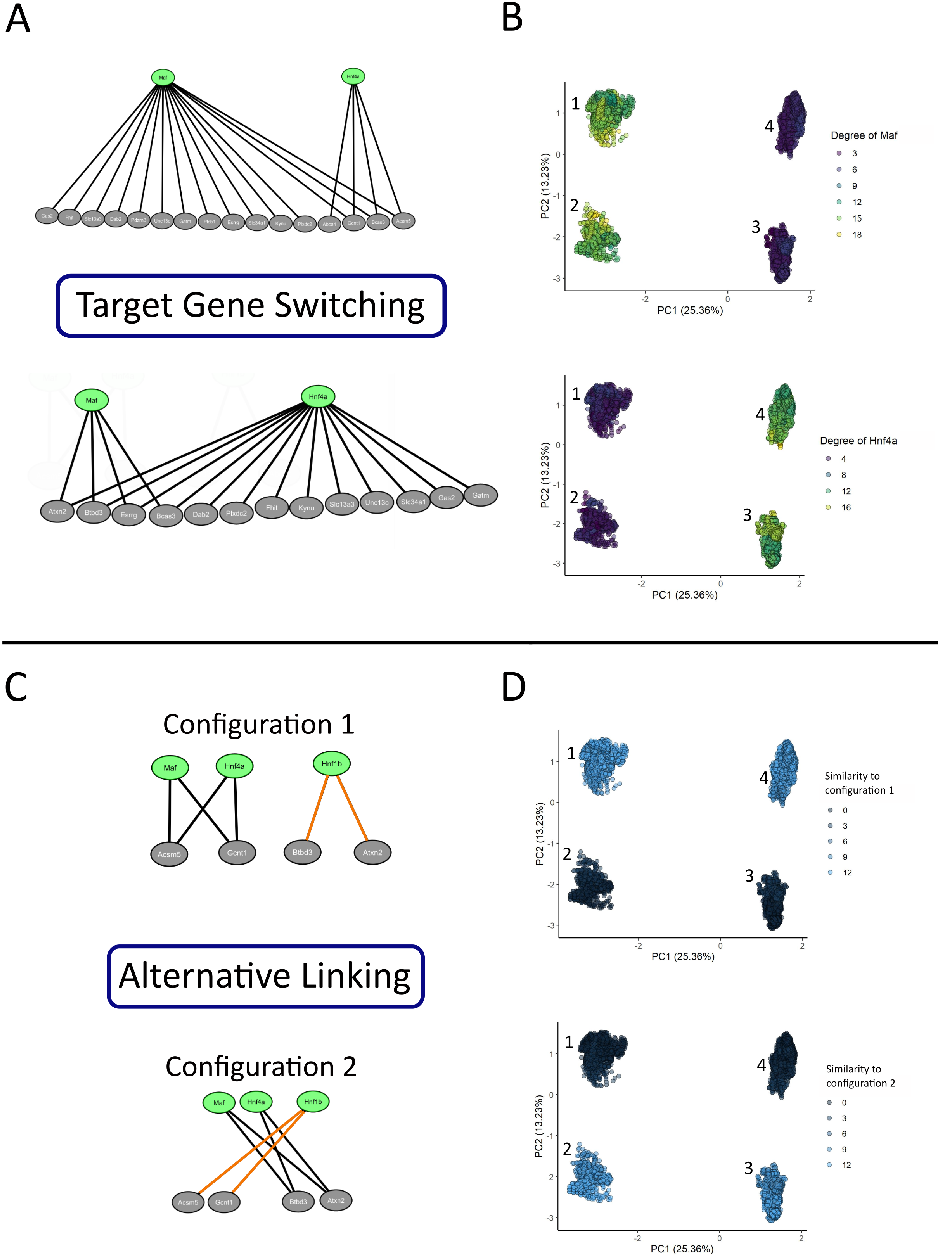
Specific network substructures discern between the different network families. (A) In clusters 1-2 Maf connects to the majority of target genes, whereas in clusters 3-4, Hnf4a connects to the majority of targets. (B) PCA feature plots coloured according to the degree of Maf (top) or Hnf4a (bottom) in each network. (C) In clusters 1 and 4, Maf and Hnf4a both typically regulate the target genes Acsm5 and Gcnt1, while Hnf1b typically regulates the targets Btbd3 and Atxn2, we label this as “configuration 1”. In clusters 2 and 3, Maf and Hnf4a typically regulate the targets Btbd3 and Atxn2, whereas Hnf1b typically regulates Acsm5 and Gcnt1, we label this as “configuration 2”. (D) PCA feature plots coloured according to the level of similarity to “configuration 1” (top) or “configuration 2” (bottom) in each network. Each configuration can be characterized by six mutually exclusive interactions, which altogether sum up to 12 interactions. Therefore, for each network solution, the similarity to configuration 1 or 2 was scored according to the existence/non-existence of these 12 interactions.

## 4 Discussion

In this study we applied RE:IN, a computational modeling tool, to model the GRN that govern kidney injury and repair. We created an ensemble of GRNs consistent with constraints from previous gene expression measurements, identified four well separated GRN families, and found the specific topological substructures that discern between them. This methodology for characterizing an ensemble of possible GRNs under a given set of experimental constraints can be generalized to other biological contexts.

Most of the contemporary tools for GRN inference such as SCENIC [1] and CellOracle [9] rely on the following information: (i) gene expression (RNA-seq) measurements; (ii) chromatin accessibility (ATAC-seq) information; and(3)known transcription factors and their motifs. However, many important regulatory mechanisms are either not well characterized yet, or involve signaling interactions and effectors that do not directly bind to cis-regulatory elements on the DNA, and thus cannot be inferred from TF motif enrichment or chromatin accessibility information. In this study we show that using a GRN inferred by SCENIC as input to RE:IN allows exploration of a large state-space of potential networks that are guaranteed to be consistent with all experimental observations. Moreover, formal reasoning provides a systematic approach to assess whether a regulatory interaction is possible or necessary for a given cell trajectory, based on a given set of experimental measurements and biochemical constraints (Figure 2).

## 5 Acknowledgments

D.M. and T.K. were supported by the Israel Science Foundation (ICORE no. 1902/12 and Grants no. 1634/13, 2017/13, and 1814/20), the Israel Ministry of Health (Grant no. 3-10146), the EU-FP7 (Marie Curie International Reintegration Grant no. 618592), the Data Science Institute at Bar-Ilan University (seed grant), the ICRF (Grant no. 19-101-PG), the Israel Ministry of Science (Grant no. 3-16220), the Israel Ministry of Justice (Veadat Haezvonot), and the Israel Cancer Association (Grant no. 20240114). The funders had no role in study design, data collection and analysis, decision to publish, or preparation of the manuscript. E.T. and H.K. were supported by the Horizon 2020 research and innovation programme for the Bio4Comp project under grant agreement number 732482 and by the ISRAEL SCIENCE FOUNDATION (Grant No. 190/19) and the Data Science Institute at Bar-Ilan University (seed grant).

## Availability of data and software code

Our models, data used to generate them and the experiments, programs, and experiments, are available at the following URL:https://github.com/TannenE/Reasoning-about-Gene-Regulatory-Network-Ensembles-in-Kidney-Injury-and-Repair. The code used to build the ABN has been described in refs. [18, 19] and made publicly available on GitHub at https://github.com/fsprojects/ReasoningEngine.

